# Unusual predominance of maintenance DNA methylation in *Spirodela polyrhiza*

**DOI:** 10.1101/2020.12.03.410332

**Authors:** Alex Harkess, Adam J. Bewick, Zefu Lu, Paul Fourounjian, Joachim Messing, Todd P. Michael, Robert J. Schmitz, Blake C. Meyers

**Affiliations:** Donald Danforth Plant Science Center, St Louis MO 63132; Department of Genetics, University of Georgia, Athens GA 30602; Waksman Institute of Microbiology, Rutgers University, New Brunswick NJ 08901; Salk Institute for Biological Studies, La Jolla CA 92037; University of Missouri – Columbia, Division of Plant Sciences, Columbia, MO 65211, USA

## Abstract

5-methylcytosine (5mC) is a modified base often described as necessary for the proper regulation of genes and transposons and for the maintenance of genome integrity in plants. However, the extent of this dogma is limited by the current phylogenetic sampling of land plant species diversity. Here, we report that a monocot plant, *Spirodela polyrhiza*, has lost CG gene body methylation, genome-wide CHH methylation, and the presence or expression of several genes in the highly conserved RNA-directed DNA methylation (RdDM) pathway. It has also lost the CHH methyltransferase *CHROMOMETHYLASE 2*. Consequently, the transcriptome is depleted of 24-nucleotide, heterochromatic, small interfering RNAs that act as guides for the deposition of 5mC to RdDM-targeted loci in all other currently sampled angiosperm genomes. Although the genome displays low levels of genome-wide 5mC primarily at LTR retrotransposons, CG maintenance methylation is still functional. In contrast, CHG methylation is weakly maintained even though H3K9me2 is present at loci dispersed throughout the euchromatin and highly enriched at regions likely demarcating pericentromeric regions. Collectively, these results illustrate that *S. polyrhiza* is maintaining CG and CHG methylation mostly at repeats in the absence of small RNAs. *S. polyrhiza* reproduces rapidly through clonal propagation in aquatic environments, which we hypothesize may enable low levels of maintenance methylation to persist in large populations.

**Significance Statement:** DNA methylation is a widespread chromatin modification that is regularly found in all plant species. By examining one aquatic duckweed species, *Spirodela polyrhiza*, we find that it has lost highly conserved genes involved in methylation of DNA at sites often associated with repetitive DNA, and within genes, however DNA methylation and heterochromatin is maintained during cell division at other sites. Consequently, small RNAs that normally guide methylation to silence repetitive DNA like retrotransposons are diminished. Despite the loss of a highly conserved methylation pathway, and the reduction of small RNAs that normally target repetitive DNA, transposons have not proliferated in the genome, perhaps due in part to the rapid, clonal growth lifestyle of duckweeds.

## Introduction

Evolutionary theory predicts that asexual populations should become less fit over time due to an irreversible accumulation of deleterious alleles (1). Duckweeds are perhaps the most striking counter-example in plants, given their cosmopolitan distribution and ability to survive in diversely harsh environments (2). Duckweed is a common name for all 36 species in the Lemnaceae family of monocots, divided across five genera: *Spirodela, Lemna, Landoltia, Wolffia*, and *Wolffiella* (**Figure 1A**). Most duckweed species rarely flower, instead reproducing primarily by rapid, clonal reproduction that occurs at one of the fastest rates in any angiosperm (3). The largest duckweed species *Spirodela polyrhiza* (~1 cm wide) intriguingly has the smallest genome size (~158 megabases) (3–5), and several genome assemblies consistently annotate fewer than 20,000 genes (6). Compared to the *Arabidopsis thaliana* genome that is roughly the same total genome size, *S. polyrhiza* has nearly 25% fewer genes. Without much meiotic recombination through sexual reproduction, and fewer genes for selection to act upon, epigenetic variation could instead be a promising mechanism to explain the global success of clonal duckweeds (7).

**Figure 1:**
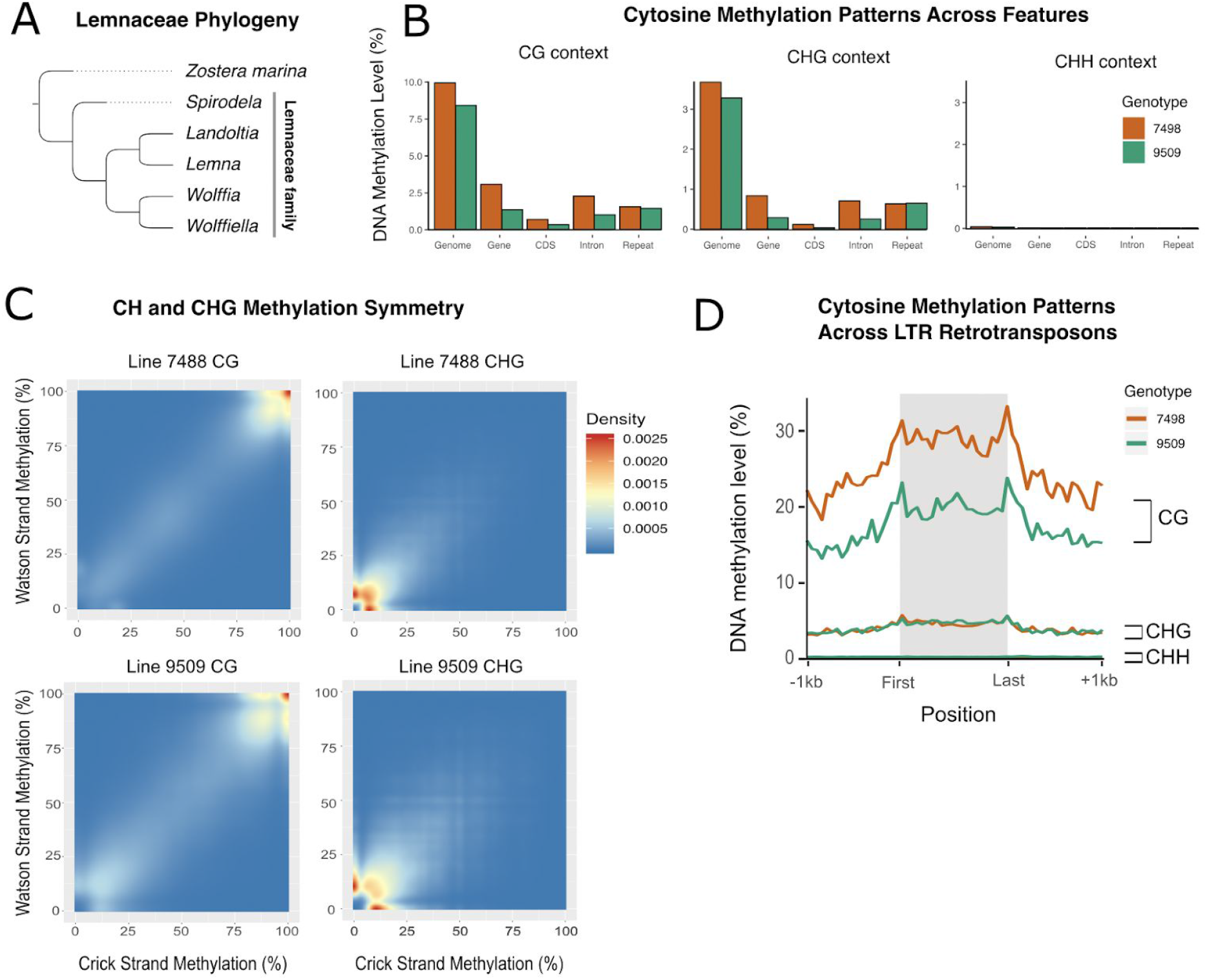
A) A condensed species tree of the Lemnaceae family, with five genera: *Spirodela, Landoltia, Lemna, Wolffia, Wolffiella*. B) DNA methylation level for whole genome, genes, CDS, introns, and repeats, across CG, CHG, and CHH site contexts, in two genotypes of *Spirodelapolyrhiza*. C) Site-wise methylation symmetry of CG and CHG on Watson and Crick strands. D) DNA methylation levels across LTR retrotransposons, across CG, CHG, and CHH sites, in two genotypes of *Spirodela polyrhiza*.

Sexual reproduction in plants is often accompanied by widespread genome-wide reinforcement of DNA methylation with localized epigenetic reprogramming in gametes (8–10). This results in patterns of both stable DNA methylation inheritance and infrequent spontaneous epialleles (11). However, nearly 60% of global crops can be bred through clonal propagation (12), highlighting the need to illustrate how epigenetics can be used to improve plant breeding efforts. Cytosine DNA methylation or 5-methylcytosine (5mC), is found in species spanning the flowering plant phylogeny (13). As the number and phylogenetic diversity of plant genomes and DNA methylomes increases, so does the observed diversity of 5mC levels, specificity and DNA methyltransferase enzymes. 5mC DNA methylation in plants occurs at three major sequence contexts, each of which require different sets of enzymes to function: CG, CHG, and CHH (where H = A, C, T). Methylation at these different contexts is established by both *de novo* and maintenance methyltransferase enzymes. DNA methylation at CG and CHG sites is typically symmetrical across the Watson and Crick strands, whereas DNA methylation at CHH sites is asymmetrical. The observed symmetry is due to the mechanisms by which 5mC is maintained after DNA replication. Methylation at CG sites relies on the maintenance methyltransferase METHYLTRANSFERASE 1 (MET1) (14–16), whereas maintenance of methylation at CHG sites relies on a positive feedback loop between dimethylation of lysine 9 on histone 3 (H3K9me2) and CHROMOMETHYLASE 3 (CMT3) (17–20). DNA methylation at CHH sites is asymmetrical and is further classified into CWA (where W = A or T) and non-CWA, based on targeting by CHROMOMETHYLASE 2 (CMT2) or by 24-nt siRNAs and DOMAINS REARRANGED METHYLTRANSFERASE 2 (DRM2), which are associated with the RNA-directed DNA methylation (RdDM) pathway, respectively (21, 22).

Variation in DNA methylation has been connected to pathogen response (23), temperature tolerance (24), and geography (25), which could be crucial attributes for clonal duckweeds given their reduced ability generate genetic variation through recombination. Intriguingly, the duckweed *S. polyrhiza* displays particularly low levels of 5mC, with evidence that low DNA methylation levels are likely related to its small genome size with the low amounts of repetitive DNA (26). However, the mechanisms underlying this variation in DNA methylation are unknown (6). Here we dissect 5mC DNA methylation patterns, histone modifications, small RNAs, and the genes that control major methylation pathways in *S. polyrhiza*. We discover that *S. polyrhiza* has lost the activity of key, canonical DNA methylation and small RNA pathway genes that consequently diminish gene body methylation, the RNA-directed DNA methylation pathway, and genome-wide CHH methylation.

## Results and Discussion

To test if low levels of 5mC might be a conserved feature across the diversity of *S. polyrhiza*, we performed whole genome bisulfite sequencing across two different genotypes (lines 7498 and 9509) (**Supplemental Table 1**). Both genotypes show similar patterns: roughly 10% of the CG sites in the genome are methylated (**Figure 1B**). Fewer than 3.28% and 3.67% of CHG and 0.0065% and 0.035% CHH sites are significantly methylated (**Figure 1B**). In *S. polyrhiza*, mCG and some mCHG (specifically CAG and CTG) are symmetrically maintained through equal DNA methylation on the Watson and Crick strands, which are normal features of maintenance methylation (**Figure 1C**). However, the maintenance of mCHG in *S. polyrhiza* is weak in comparison to other species that possess a functional CMT3 (13). CG methylation is present at small clusters of Long Terminal Repeat (LTR) retrotransposons in the genome (**Figure 1D**), but CHH methylation which is normally enriched in repetitive elements like LTR retrotransposons (27), is absent (**Figure 1C-D**).

DNA methylation at some CHH sites is guided by small RNAs (sRNAs) generated through the RdDM pathway (28) (**Figure 2A**), so small RNA (sRNA) sequence reads were generated for both *S. polyrhiza* lines to test for functional defects in the pathway (**Supplemental Table 2**). Both lines display a distinct lack of 24-nucleotide (nt), heterochromatic siRNAs (het-siRNAs) which are typically the most abundant size class of angiosperm sRNAs (29, 30) (**Figure 2B**). The highly conserved, canonical RdDM pathway produces these 24 nt het-siRNAs via DICER-LIKE 3 (DCL3) processing of an RNA Polymerase IV (Pol IV)-derived double-stranded RNA (31, 32). These DCL3-derived sRNAs are loaded into ARGONAUTE 4 (AGO4) and guided to their sites of action (**Figure 2A**) (28, 33, 34). Due to the reduction of 24 nt het-siRNAs, available whole plant mRNA-seq data was mined for evidence of the expression of RdDM-related genes (**Figure 2C**). *DCL3* is present in the genome as a seemingly full-length sequence with no in-frame stop codons, but no gene expression was detectable (FPKM <1). *DCL3* expression is also not detected under various growth and stress conditions in *S. polyrhiza*, including copper, kinetin, nitrate and sucrose additions (35) (**Supplemental Fig. 1**). The *DCL3* upstream region is short (fewer than 200 nt), and possibly interrupted by another gene, which may entirely disrupt *DCL3* gene activity (**Supplemental Fig. 2**).

**Figure 2:**
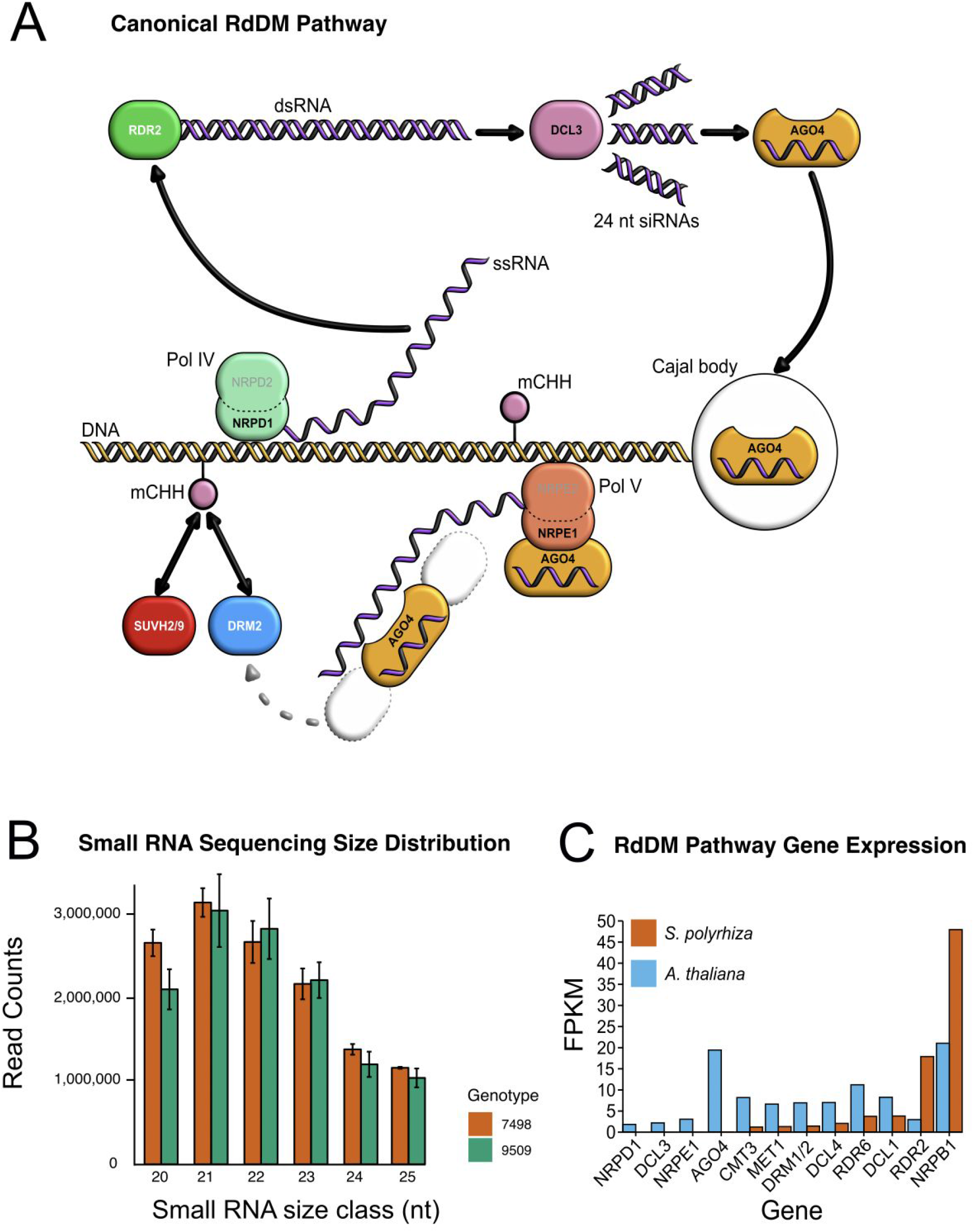
A) Diagram of the canonical RNA-directed DNA methylation (RdDM) pathway in plants. RNA Polymerase IV (Pol IV) transcribes a single-stranded RNA (ssRNA) which is converted to a double-stranded RNA (dsRNA) by RNA-DIRECTED RNA POLYMERASE 2 (RDR2). DICER-LIKE 3 (DCL3) then cleaves those dsRNA products into 24 nucleotide small RNA (sRNA) products. One strand of each sRNA is loaded into ARGONAUTE 4 (AGO4), and the AGO-sRNA complex binds to complementary RNA sequences transcribed by RNA Polymerase V (Pol V), guided by interaction with SUVH2 and SUVH9. DOMAINS REARRANGED METHYLTRANSFERASE 2 (DRM2) is then recruited, which guides methylation of DNA at those sites. B) The distribution of small RNA sequence read abundance between 20-25 nucleotides in two genotypes of *S. polyrhiza*. C) Gene expression in *S. polyrhiza* line 9509 and *A. thaliana* measured by RNA-seq for several RdDM and methylation-related genes.

Given an absence of detectable *DCL3* expression (**Figure 2C**), we investigated the presence and expression of orthologs of other plant Dicer-like genes (*DCL1, DCL2, DCL4*). *DCL1*, which functions in microRNA (miRNA) production, is expressed and produces many conserved miRNAs, indicating it functions normally (36). However, *DCL2*, which functions largely in viral defense, is missing from the *S. polyrhiza* genome (37) (**Supplemental Fig. 3**). *DCL4*, which generates 21-nt siRNAs, is present in the genome and expressed (38–40). *DCL5*, which is implicated in phased siRNA production in maize (41) and has a role in flower fertility (42), is also not present in the genome (30) (**Supplemental Fig. 3**). In addition to *DCL2* and *DCL3*, there was no detectable expression for *AGO4*, nor the genes encoding the two major catalytic subunit genes of the Pol IV complex *(NRPD1* and *NRPE?*) that transcribe single-stranded RNA precursors from RdDM regions and are required for siRNA and methylation-dependent heterochromatin formation (43) (**Figure 2C**). *CMT3* and *MET1* are expressed in *S. polyrhiza*, consistent with their roles in the maintenance of CG and CHG methylation in the *A. thaliana* genome (**Figure 2C**). We next tested whether the lack of expression of some RdDM genes is a conserved phenomenon across some or all duckweed species in the Lemnaceae family. *De novo* transcriptome assemblies of publicly available whole plant RNA-seq data for species from two genera of duckweeds, *Landoltia punctata* and *Lemna minor* (**Figure 1A**), were interrogated for Dicer-like gene expression. In both *L. punctata* and *L. minor, de novo* transcripts were assembled for *DCL1* and *DCL4*, however there were no assemblies with BLASTX hits (1e-10) to *DCL2* and *DCL3*. Although whole genome assemblies of species representing all five genera of duckweeds will be needed to definitively test this hypothesis, these data suggest that the expression loss of *DCL2* and *DCL3*, possibly leading to the loss of canonical RdDM, may be a widespread phenomenon across several genera of the Lemnaceae family (**Figure 1A**).

Although RdDM is one route to forming CHH methylation, an RdDM-independent mechanism is through the action of CHROMOMETHYLASE 2 (CMT2), a plant-specific DNA methyltransferase that is highly conserved across angiosperms (21, 44, 45) (**Figure 3A**). CHH sites targeted by the RdDM pathway typically show enrichment at all contexts (21, 46), which *S. polyrhiza* does not exhibit (**Figure 3A**). In *A. thaliana*, CHH methylation deposited via CMT2 can be distinguished from RdDM-targeted sites given that they show an enrichment of CWA methylation (W = A or T) relative to other contexts (21, 46) and they are enriched at regions possessing H3K9me2 (44). However, a *CMT2* homolog is missing from the *S. polyrhiza* genome (lines 7498 and 9509) (**Figure 3B**). As expected given the loss of *CMT2*, there is no enrichment of CWA methylation in either genotype (**Figure 3C**). There is a low level of CWG methylation in both lines, though (**Figure 3C**). CWG methylation is dependent on *CMT3* (21), which is present and expressed (**Figure 2C**). Across the global range of *A. thaliana*, there is extensive variation at the *CMT2* locus including a non-functional *cmt2* allele that is associated with reduced genome-wide CHH methylation, but also the benefit of an increased tolerance to heat stress (24, 47). Given that *S. polyrhiza* is globally distributed and thrives in a variety of climates and stresses, increased genotyping and phenotyping of diverse populations may reveal similar patterns of methylation-sensitive phenotypes. Intriguingly, *CMT2* is missing in the maize genome (48), but also missing from the aquatic seagrass *Zostera marina* genome assemblies and annotations (**Figure 3B**), suggesting that *CMT2* loss may be a shared feature that has evolved in multiple aquatic plants in the Alismatales order. Despite a lack of expression of key RdDM genes and sRNAs that normally function to target repetitive DNA, there has not been a recent detectable expansion of LTR retrotransposons in the *S. polyrhiza* genomes (**Figure 3D**) (3, 26), nor are they methylated in the typical CHH context (**Figure 1D**). Specifically, only 3/1114 (0.003%) and 6/1510 (0.004%) LTR retrotransposons are enriched for CHH methylation in 7498 and 9509 genomes, respectively, and likely false positives (48).

**Figure 3:**
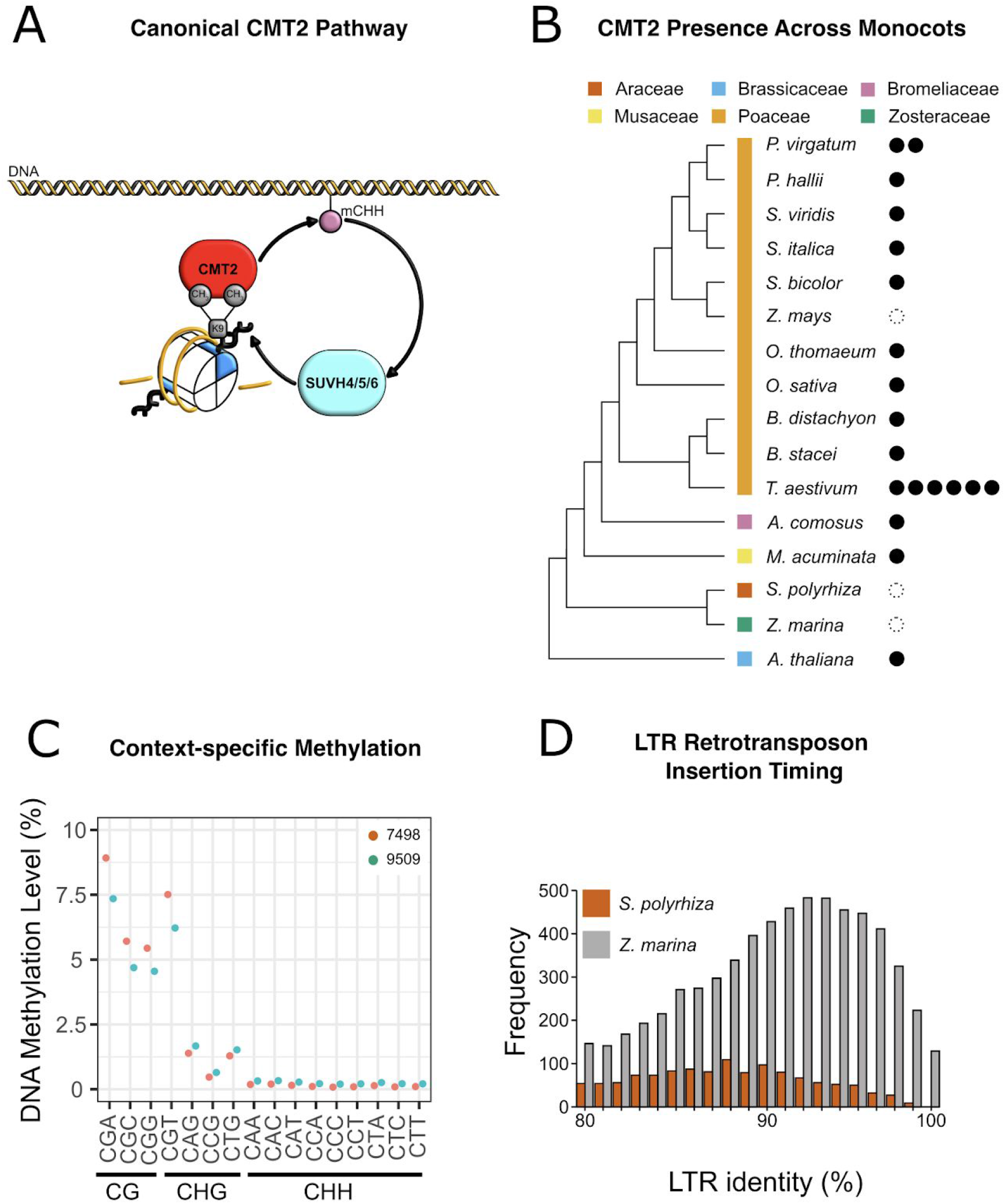
A) Diagram of the canonical CHROMOMETHYLASE 2 (CMT2) pathway. B) Presence (filled circles) and absence (empty circles) of CMT2 homologs in genomes across the monocots. C) Genome-wide DNA methylation of two genotypes of *S. polyrhiza* split into all possible contexts. D) Relative LTR retrotransposon insertion timings between *S. polyrhiza* and *Zostera marina*, based on LTR percent identity comparisons.

The loss of CHH methylation, 24 nt het-siRNAs, and CMT2 suggests that the abundance of heterochromatin may also be low. Cao et al. (49) made an observation using 5mC and histone 3 lysine 9 di-methylated (H3K9me2) immunostaining, a common histone modification in heterochromatic regions of the genome (50), that *S. polyrhiza* and four other genera of the Lemnaceae lack strong signals of concentrated heterochromatic blocks of DNA. H3K9 methylation mediates CHG and CHH methylation through the action of CMT3 and CMT2, respectively (44). To test if the CMT2 loss and the weak levels of CHG methylation is tied to a reduction of H3K9 methylation in *S. polyrhiza*, we performed chromatin immuno-precipitation sequencing (ChIP-seq) of H3K9me2 (**Supplemental Table 3**). H3K9me2 is sparsely distributed throughout the euchromatic chromosome arms and shows a discrete enrichment of a large domain within each chromosome (**Fig 4A**). These relatively larger domains of H3K9me2 are approximately 400-600 kb and presumably reflect the pericentromeric regions similar to observations in other angiosperms like *A. thaliana* (**Fig 4A**). H3K9me2 occupies ~15% of the line 9509 genome. In the 9509 genome, 746/1,510 (49.40%) LTR retrotransposon annotations overlap H3K9me2 (Fisher’s Exact Test, p < 0.001; **Supplemental Table 4**, **Supplemental Fig, 4**). Overall, H3K9me2 and heterochromatin appears normal in *S. polyrhiza*, especially when considering the small genome size split into 20 chromosomes.

**Figure 4:**
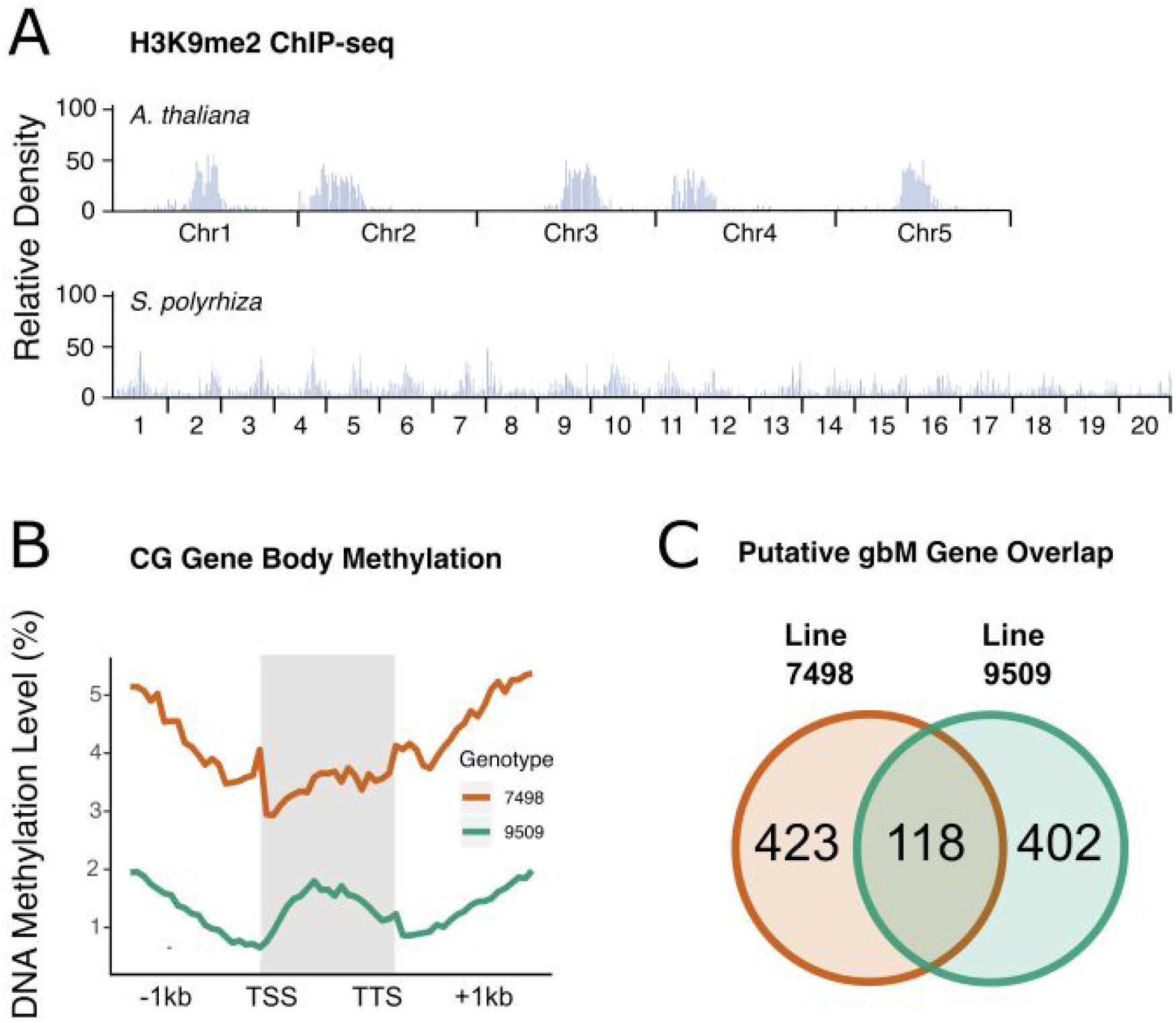
A) Distribution of H3K9me2 ChIP-seq peaks in *A. thaliana* and *S. polyrhiza* line 9509. B) Weighted gene body methylation plotted along CDS regions spanning from Transcription Start Site (TSS) to Transcription Termination Site (TTS), plus or minus 1 kilobase. C) The overlap of blindly calling putative gene body methylated genes in two genotypes of *S. polyrhiza*.

Maintenance of DNA methylation at heterochromatin is associated with the establishment and maintenance of gene body DNA methylation (gbM) (46, 51–53). It is characterized by an enrichment of CG DNA methylation between the transcription start site and transcription termination site of genes (45, 54). Genes with gbM are typically moderately expressed throughout all tissues, long and exhibit low rates of nucleotide substitutions compared to non-gbM genes (55, 56). We blindly quantified CG methylation in coding regions of each gene, only accounting for the number of methylated CG sites, total CG sites, and read coverage (**Figure 4C**). In *S. polyrhiza*, this resulted in 541 and 520 putative gbM genes in lines 7498 and 9509, respectively, or 2-3% of the total gene annotation set. Comparing the two putative gbM gene sets, 118 genes overlap between the two genomes (blastp 1e-40), which is unexpected as gbM genes are often highly conserved (53, 55) (**Figure 4B**). These results are similar to another species that has lost gbM, *Eutrema salsugineum*, where roughly 500 genes were bioinformatically detected as having gbM signatures using similar methodology (52). This result is likely driven by a similar false positive rate of gbM gene detection in both species, as well as transposon misannotation, and that like *E. salsugineum*, gbM has been lost in *S. polyrhiza*.

The faithful establishment and maintenance of gbM is tied to a self-reinforcing feedback loop that relies on the interplay between CMT3 and H3K9me2 (17, 46, 53, 57). This is further supported by studies in *A. thaliana* whereby mutants that result in a loss of maintenance of heterochromatin lead to ectopic activity of CMT3 in gbM genes (58–60). In *S. polyrhiza*, we show that the CMT3/H3K9me2 feedback loop is weak in comparison to other angiosperms, even though H3K9me2 has a typical distribution throughout the genome. Therefore, it is possible that CMT3 activity is impaired, which leads to a weakly functioning feedback loop in *S. polyrhiza* and the loss of gbM. These results are consistent with proposed models from Wendte et al. (46) and Inigaki and Kakutani (51), in which CMT3 and H3K9me2 work coordinately to establish *de novo* gbM.

Ecological life history and developmental traits may strongly influence genome-wide patterns of DNA methylation and inheritance, especially relating to the suppression of transposon expansion over time. *S. polyrhiza* primarily reproduces via rapid clonal propagation rather than by flower production and sex (61), though some low frequency instances of flowering have been reported (62, 63). The methylomes of other clonally propagated species *(Eucalyptus grandis, Fragaria vesca, Manihot esculenta, Theobroma cacao*, and *Vitis vinifera)* possess mCHH, although the levels are lower than non-clonally reproducing angiosperms (13, 64). This suggests that CHH reinforcement is linked to sexual reproduction, but isn’t necessary for transposon silencing as clonally propagated species rely more on maintenance DNA methylation. Resequencing of globally distributed *S. polyrhiza* accessions reveals very little per-site genetic diversity within the species, a low recombination rate, and weak purifying selection, but still a large effective population size (*N_e_*) (65, 66). Few transposons exist in the ~150 Mb *S. polyrhiza* genome, but a high ratio of solo-LTRs to intact LTR retrotransposons (26) suggests that LTR excision is actively occurring despite weak purifying selection. Individuals with fewer transposons and the ability to excise them may have selective advantages in large populations (67), given that a deleterious transposon insertion is unlikely to propagate and fix in a large clonal population, possibly compensating for the lack of a canonical RdDM pathway.

Several alternative hypotheses could explain the lack of CHH methylation and 24 nt het-siRNAs. DNA methylation and sRNA production may be limited to cell or tissue-specific regions, such as the developing meristematic region where daughter plantlets emerge. Tissue sampling for duckweeds is often performed on several individual plantlets combined, given the small size, which would reduce the detectable signal of tissue- or cell type-specific changes and require more precise excision or cell sorting techniques. Overall, *S. polyrhiza* displays a loss of CHH-type DNA methylation and heterochromatic siRNAs, which may be tied to its rapid asexual reproduction. Our work in *S. polyrhiza* demonstrates that reproductive success through rapid clonal propagation may benefit from the sacrifice of the RdDM and CMT2 pathways.

### Data Availability

All raw small RNA, DNA methylation, and H3K9me2 ChIP-seq data is available at BioProject GSE161234. Epigenome browsers are available for *S. polyrhiza* line 7498 (http://epigenome.genetics.uga.edu/SchmitzLab-JBrowse/?data=spi_pol_7498) and for line 9509 (http://epigenome.genetics.uga.edu/SchmitzLab-JBrowse/?data=spi_pol_9509).

## Author contributions

The study was conceived by AH, AJB, RJS, and BCM. AH and AJB performed genomic, phylogenetic and evolutionary analyses. AH, AJB, TM and PF performed gene expression and sRNA analyses, and AJB performed DNA methylation analyses. ZL generated and analyzed H3K9 ChIP data. PF and JM provided materials and assistance with expression analyses. AH and AJB wrote the article with contributions made by all authors.

## Materials and Methods

### DNA methylation sequencing and sequence alignment

Whole-genome bisulfite sequencing data were generated according to (68). Single-end short read libraries (150 bp) were aligned using the methylpy pipeline (69) to the *S. polyrhiza* 7498 and 9509 genomes. Methylpy calls programs for read processing and aligning: (i) reads were trimmed of sequencing adapters using Cutadapt (70), (ii) and then mapped to both a converted forward strand (cytosines to thymines) and converted reverse strand (guanines to adenines) using bowtie (71). Reads that mapped to multiple locations, and clonal reads were removed. The chloroplast genome (GenBank: JN160603.2) was used to estimate a rate of sodium bisulfite non-conversion.

### DNA methylation analyses

DNA methylation levels were estimated as weighted DNA methylation, which is the total number of aligned DNA methylated reads divided by the total number of methylated plus un-methylated reads with a minimum coverage of at least 5 reads (72). Global weighted DNA methylation was estimated across the entire genome, within intergenic regions, transposons, genes (exons+introns), exons and introns. Additionally, the genome was divided into non-overlapping 50000 bp windows, and weighted DNA methylation was estimated for each window.

For metaplots, the locus body – start-to-stop codon for genes and first to last bp for transposons – was divided into 20 proportional windows based on locus length. Within gene bodies only sequenced reads mapping to coding, exonic DNA were used. Additionally, 1000 bp upstream and downstream were divided into 20 proportional windows. A single weighted DNA methylation value was calculated for each window across all loci.

For each gene, a binomial test with a Benjamini–Hochberg False Discovery Rate (FDR) correction was applied to determine enrichment of DNA methylation at the three sequence contexts (CG, CHG, and CHH). Only CG, CHG and CHH sites found within coding, exonic sequences were considered. The weighted DNA methylation level of cytosines at CG, CHG and CHH sites across all coding regions were used as the probability of success, respectively. Enrichment tests for gene body methylation were performed using code from (53), found at https://github.com/schmitzlab/Natural_variation_in_DNA_methylation_homeostasis_and_the_emergence_of_epialleles.

To determine per-site methylation levels, the weighted DNA methylation level for each cytosine with ≥ 3 reads of coverage was calculated. Additionally, DNA methylation levels of symmetrical cytosines (CG or CWG, W = A|T) with ≥ 3 sequencing coverage were estimated for each strand (Watson and Crick). All plots were generated in R v3.2.4 (https://www.r-project.org/).

### sRNA and mRNA sequencing analysis

sRNA sequencing reads were generated using whole plant total RNA isolated using TRI reagent and the Somagenics RealSeq-AC kit with 100 ng of total RNA as input. Reads were adapter-trimmed with cutadapt v2.0 (70) with options “-m 15 TGGAATTCTCGGGTGCCAAGG”. Cleaned reads were aligned to the reference genome using bowtie with settings “-a -v 0” to only report end-to-end alignments with zero mismatches.

Raw mRNA-Seq reads from strain 9509 were retrieved from the Sequence Read Archive (SRR3090696), cleaned with Trimmomatic v0.32 with settings “ILLUMINACLIP:2:30:10 LEADING:3 TRAILING:3 SLIDINGWINDOW:4:15 MINLEN:50” and aligned to the reference genome with TopHat v2.1.1 with default settings other than “-i 25”. Per-gene expression was calculated with Cufflinks v2.2.1 with default settings.

### Chromatin immunoprecipitation sequencing (ChIP-seq) and analysis

ChIP was performed as previously described (73). Briefly, 1 g of fresh duckweed plantlets were crosslinked in 1% formaldehyde for 10 min. Nuclei were then isolated and sonicated for 15 min, twice. Histone-DNA complexes were pulled down with anti-H3K9me2 (Cell Signaling Technology antibody #9753s). DNA was isolated and used to prepare ChIP-seq libraries with the TruSeq ChIP Library Preparation Kit (Illumina, IP-202-1012). Sequencing was performed on an Illumina NextSeq500 in Georgia Genomics and Bioinformatics Core (GGBC) in the University of Georgia.

Raw ChIP reads were trimmed for adapters and low-quality bases using Trimmomatic with the following options: reads were trimmed for TruSeq version 3 single-end adapters with a maximum of two seed mismatches, palindrome clip threshold of 30, and simple clip threshold of 10. Trimmed reads were mapped to the genome using bowtie1 with “-v 2 --best --strata -m 1” (71). Mapped reads were sorted using SAMtools (74) and then clonal duplicates were removed using picard (http://broadinstitute.github.io/picard/). Remaining reads were converted to BED format with Bedtools (75). H3K9me2 enriched regions were identified with MACS2 with parameter “--keep-dup all --broad” (76). Enrichment of H3K9me2 overlaps with LTR retrotransposons was tested using a Fisher’s Exact Test implemented in bedtools v2.26.0.

### Phylogenetic analyses

CHROMOMETHYLASE (CMT) protein sequences were obtained from (45), and additional sequences were identified in monocot species listed on Phytozome v12 (https://phytozome.jgi.doe.gov/pz/portal.html) using best blastp hit e-value ≤ 1E-06 and bit score ≥ 200) to *A. thaliana* CMT1 (AT1G80740.1), CMT2 (AT4G19020.1) and CMT3 (AT1G69770.1). Similarly, DICER-LIKE (DCL) homologs were identified in all monocot species listed on Phytozome v12 using best BLASTP hit to *A. thaliana* DCL1 (AT1G01040.2), DCL2 (AT3G03300.1), DCL3 (AT3G43920.2), and DCL4 (AT5G20320.1). Protein sequences were aligned using the program PASTA with default parameters. Following alignment, GBblocks was used to identify conserved amino acid positions. All parameters were kept at the default setting except −b2=*n*0.66 where *n* is the number of sequences and −b5=h. BEAST v2.3.2 was used to estimate the phylogeny with a BLOSUM62 substitution matrix. The MCMC chain in BEAST was allowed to run until stationarity and convergence (ESS≥200) was reached, and was assessed using the program Tracer v1.6. A maximum clade credibility tree was generated from the posterior distribution of trees with the burn-in removed using the program TreeAnnotator v2.3.2. Finally, the program FigTree (http://tree.bio.ed.ac.uk/software/figtree/) was used to visualize the tree and exported for stylization. Alignment, site filtering, and tree estimation was performed identically and separately for CMT and DCL phylogenies.

### Comparative transcriptome analyses

To estimate the phylogenetic placement of the loss of DCL2 and DCL3 expression, SRA RNA-seq data were downloaded for *Landoltia punctata* (SRR647050/ and *Lemna minor* (SRR2917879). Data were cleaned and assembled using Trinity v2.5.1 with default options. Assemblies were subject to blastx searches (1e-10) against the present, but not expressed *Spirodela* DCL3 gene model annotation predicted peptide (Spipo14G0010100).

### LTR Retrotransposon annotation

LTR retrotransposons were annotated *de novo* using GenomeTools LTRharvest with options “-similar 85 -mindistltr 1000 -maxdistltr 15000 -mintsd 5 -maxtsd 20”.

## Acknowledgements

We thank the Georgia Advanced Computing Research Center (GACRC) for computational resources. We are grateful to Yinwen Zhang for assistance in interpreting gene body methylation code, and to Brigitte T. Hofmeister for creating and maintaining the browsers. This work was funded by an NSF National Plant Genome Initiative Postdoctoral Fellowship to AH (#1611853). This study was supported by the National Science Foundation (MCB-1856143) to RJS.

