## Supplemental Material for "Unusual predominance of maintenance DNA methylation in *Spirodela polyrhiza*"

**Fig. S1.**

*DCL3* is not expressed under various stress conditions using aligned RNA-seq data of several stresses, including copper, kinetin, nitrate, and sucrose additions.

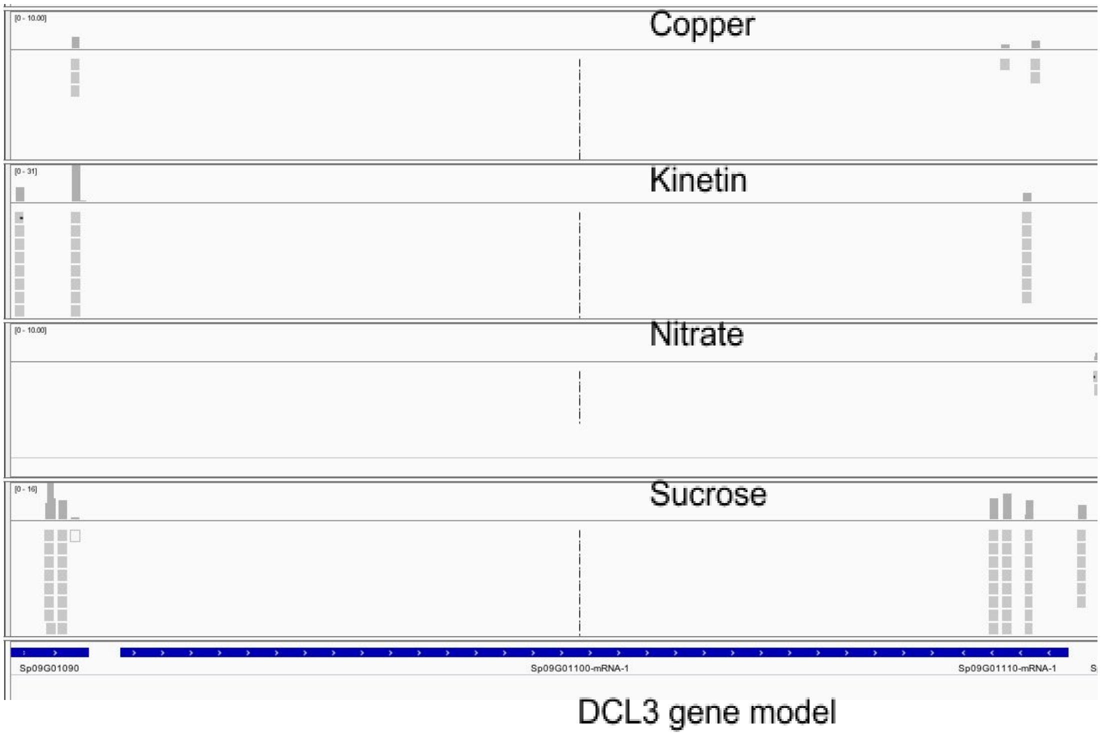

**Fig. S2.**

A) DCL3 protein structure with canonical Dicer-like PFAM domains. DEAD/DEADH-box helicase, Helicase C-terminal domain, Dicer dimer, PAZ, and two tandem Ribonuclease III domains are intact in the gene model when the protein was screened with hmmscan against the PFAM database. B) The *DCL3* gene model is not expressed when measured with RNA-seq read coverage, and it has a short promoter region (highlighted in red) that is possibly interrupted by the upstream U-box protein SP09G01090.

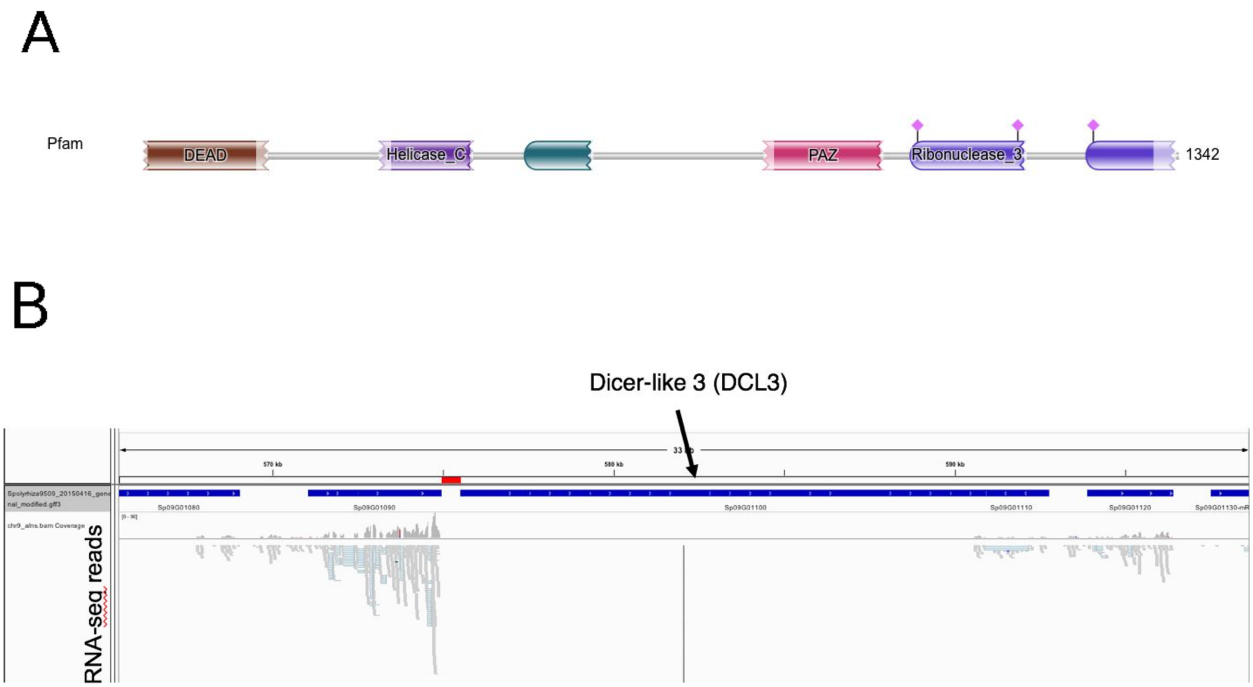

**Fig. S3.**

A) Dicer-like protein family phylogenetic tree (DCL1, DCL2, DCL3, DCL4, DCL5) and B) ZMET/CMT Bayesian phylogenetic tree built using BEAST. Nodes are annotated with a ball size proportional to a posterior probability (see the key in each panel).

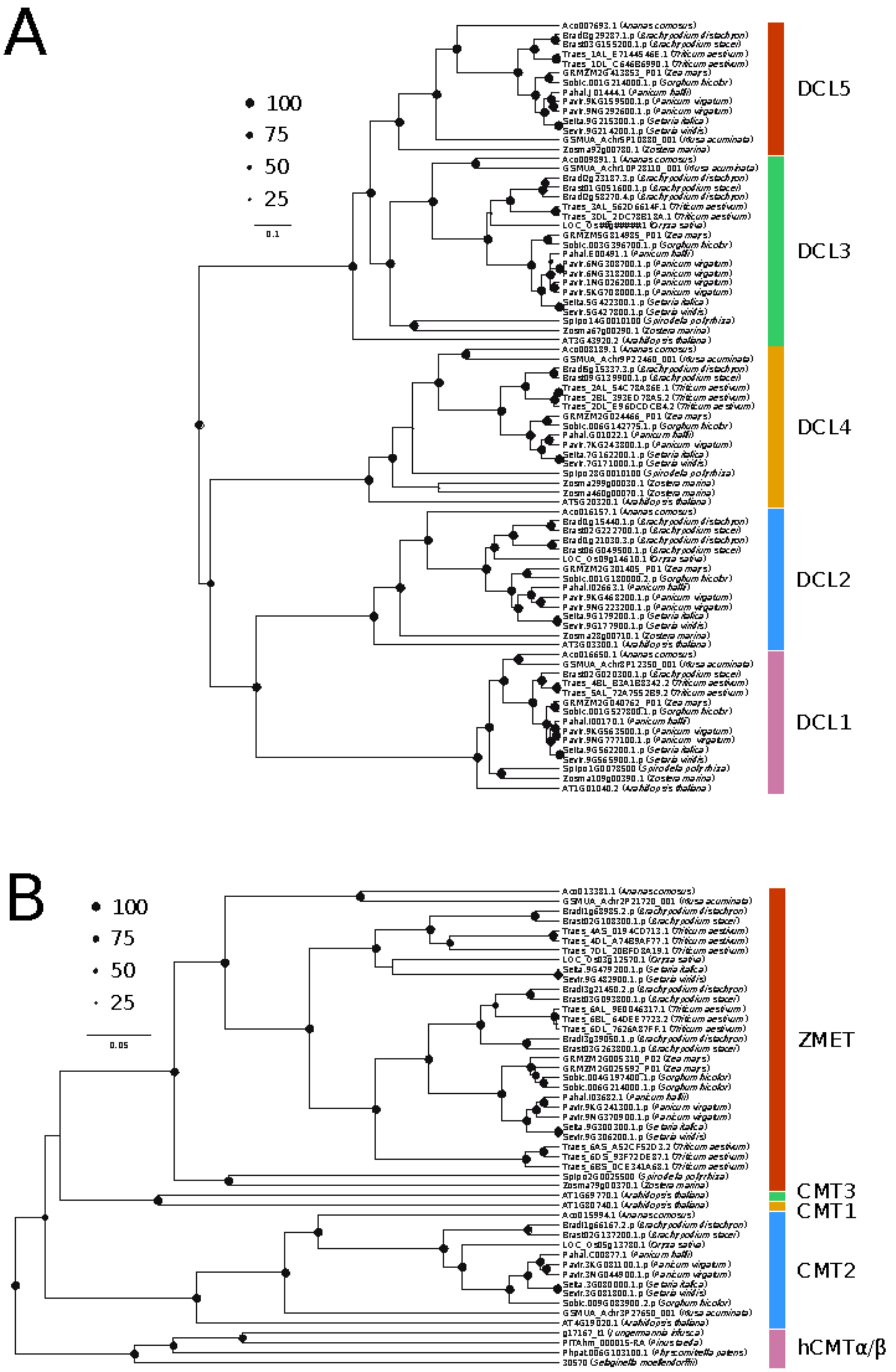

**Fig. S4: Enrichment of H3K9me2 at LTR retrotransposons**

**H3K9me2 enrichment at LTR retrotransposons** (Sp9509 Chromosome 1)

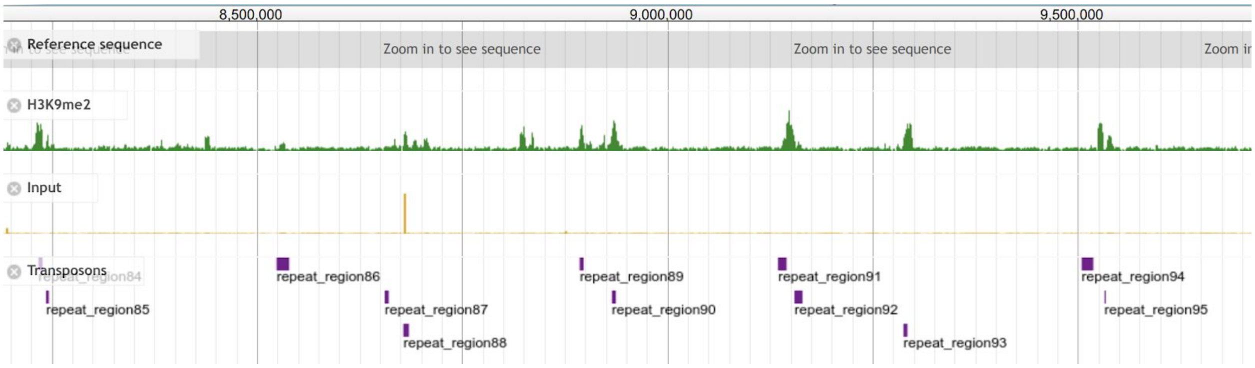

**Table S1.**

Sequencing and alignment statistics for *S. polyrhiza* line 7498 and 9509 MethylC-seq

| Library | Raw Reads | Alignments to genome after PCR duplicate removal | Non-Conversion rate (%) |
|---|---|---|---|
| Sp9509_methylC | 113,081,904 | 46,730,451 | 1.65 |
| Sp7498_methylC | 58,068,357 | 32,028,794 | 0.13 |

**Table S2.**

Sequencing and alignment statistics for *S. polyrhiza* line 7498 and 9509 small RNA data.

| Library | Raw Reads | Adapter and 18-30 nt length trimmed reads | Perfect alignments to genome |
|---|---|---|---|
| 7498_rep1 | 64,619,776 | 46,257,454 | 23,346,108 |
| 7498_rep2 | 67,122,721 | 51,974,074 | 24,270,971 |
| 9509_rep1 | 74,825,605 | 66,368,896 | 55,685,485 |
| 9509_rep2 | 73,778,606 | 66,612,352 | 48,810,499 |

**Table S3.**

Sequencing and alignment statistics for *S. polyrhiza* line 7498 and 9509 H3K9me2 ChIP-Seq

| Library | Raw Reads | Alignments to genome after PCR duplicate removal |
|---|---|---|
| Sp7498_H3K9me2 | 30,509,305 | 26,286,244 |
| Sp7498_input | 45,911,383 | 37,887,965 |
| Sp9509_H3K9me2 | 24,523,030 | 23,607,285 |
| Sp9509_input | 82,794,333 | 60,816,027 |

**Table S4.**

Fisher's Exact Test overlap of LTR retrotransposons and H3K9me2 peaks

| | Overlapping an LTR | Not overlapping an LTR |
|---|---|---|
| Overlapping an H3K9me2 peak | 713 | 508 |
| Not overlapping an H3K9me2 peak | 1,756 | |

*Two-tailed Fisher's Exact P-value = 2.0654e-184
